# An updated and extended atlas for corresponding brain activation during task and rest

**DOI:** 10.1101/2020.04.01.020644

**Authors:** Marlene Tahedl, Jens V. Schwarzbach

## Abstract

The complexity of our actions and thinking is likely reflected in functional brain networks. Independent component analysis (ICA) is a popular data-driven method to compute group differences between such networks. To aid interpretation of functional network analyses, Smith and colleagues proposed a template of ten functional networks identified in 36 healthy participants. They labeled them those components according to their similarity with statistical parameter maps from a metaanalysis of task-based functional magnetic resonance imaging studies (Smith et al., 2009). However, those original templates show substantial distortion with respect to what up-to date correction methods can achieve, such that trying to capture relevant effects within several cortical regions, especially the sensorimotor and orbitofrontal cortices, as well as the cerebellum may yield suboptimal results. Here, we provide a technical update and extension to the original templates. Using correlation analyses, we identified the best matching maps of each of the original ten templates to ICA maps from the Human Connectome Project (HCP). The HCP provides group-parcellations of a large dataset (n = 1003) with high-quality data. This approach yields a better fit of spatial component maps with anatomical borders and gray-/white-matter-boundaries. Additionally, we provide a version of the updated templates in CIFTI file-format, an emerging format in the neuroimaging community that combines surface-based data with subcortical/cerebellar data in volumetric space. The two formats we provide here offer an improvement on the templates by Smith et al., which should enhance sensitivity and interpretability of future research that compares functional networks between groups.

## Introduction

Functional magnetic resonance imaging at rest (resting-state fMRI or rs-fMRI) has revealed consistent patterns of statistical associations, so-called networks of functional connectivity, between certain sets of regions in the human brain (Biswal, Yetkin, Haughton, Hyde, 1995; Fox Raichle, 2007; Smith et al., 2009). In this context (spatial) independent component analysis (ICA) has become a widely adopted multivariate data driven analysis method, which attempts to identify spatially independent components (i.e. statistical parameter maps) whose members share a common time-course (Calhoun, Liu, Adali, 2009; Hyvärinen, 1999). In the last wo decades, studies using rs-fMRI have reported a consistent set of large-scale resting state networks whose functions have become research topics in cognitive and clinical neurosciences (Damoiseaux et al., 2006; Raichle et al., 2001; Yeo, Krienen, Sepulcre, Sabuncu, others, 2011). In order to compare ICA components from rs-fMRI on the group level (e.g. patients vs. controls) there exist two common strategies, both of which rely on the so-called dual-regression approach (Beckmann, Mackay, Filippini, Smith, 2009; Nickerson, Smith, Öngür, Beckmann, 2017). In the first approach, one produces an ICA-group map on the concatenated data of all subjects, yielding a set group maps and corresponding time courses (step 1 in dual regression). From these, subject specific maps are calculated (step 2 in dual regression), which serve as the basis for group comparisons. The advantage of this approach is that the group maps are optimized for the sample of subjects in the study. However, since the results of group-ICA decompositions can look different depending on acquisition parameters, groups sice, preprocessing strategies, it may become difficult to compare the results of different studies. The alternative approach to computing the ICA group map from the study sample (step 1 in dual regression) is to use a common template with one map per component of interest. These maps can originate from a particular study or for example from a common template. In a seminal paper based on 36 healthy participants, Smith and colleagues (2009) pointed out a striking similarity in the spatial structure between commonly reported resting-state maps and task-based statistical parameter maps drawn from meta-analyses of 1687 fMRI studies that investigated different perceptual, cognitive, and motor-tasks in overall 29671 subjects. They have made this set of maps available as a template (https://www.fmrib.ox.ac.uk/datasets/brainmap+rsns/), which has been widely used either for providing a standardized visualization of results or as the basis for computing individual component maps (Castellazzi et al., 2018; Pflanz et al., 2015; Rane et al., 2014; Rubin et al., 2017).

Computing individual maps based on a common template, is highly interesting for the cognitive and clinical neurosciences in terms of standardization, and it requires that these maps are as accurate as possible in order to yield high sensitivity and specificity. Detailed inspection of the template reported by Smith and colleagues shows that these templates suffer from spatial distortions such that the contain voxels outside gray-matter, but even more importantly that some brain structures such as motor cortex, (orbito-) frontal cortex, and cerebellum are not well covered (see Figs 1, 4). This reduces specificity and sensitivity of approaches that use this template in the first stage of a dual regression analysis for creating individual ICA maps for the second stage. Here, we propose to make use of the high-resolution ICA-maps from the Human Connectome Project 1200 Subjects Release (www.humanconnectome.org) (Marcus et al., 2011; Van Essen et al., 2013) to create an updated template that provides more accurate and complete brain coverage while keeping the functional topology as similar as possible to the original maps. Furthermore, we provide the template also in CIFTI (Connectivity Informatics Technology Initiative, https://www.nitrc.org/projects/cifti/) file-format in so-called “grayordinates”, combining cortical data rendered on the surface with volumetric data of subcortical structures and the cerebellum.

**Fig. 1.**
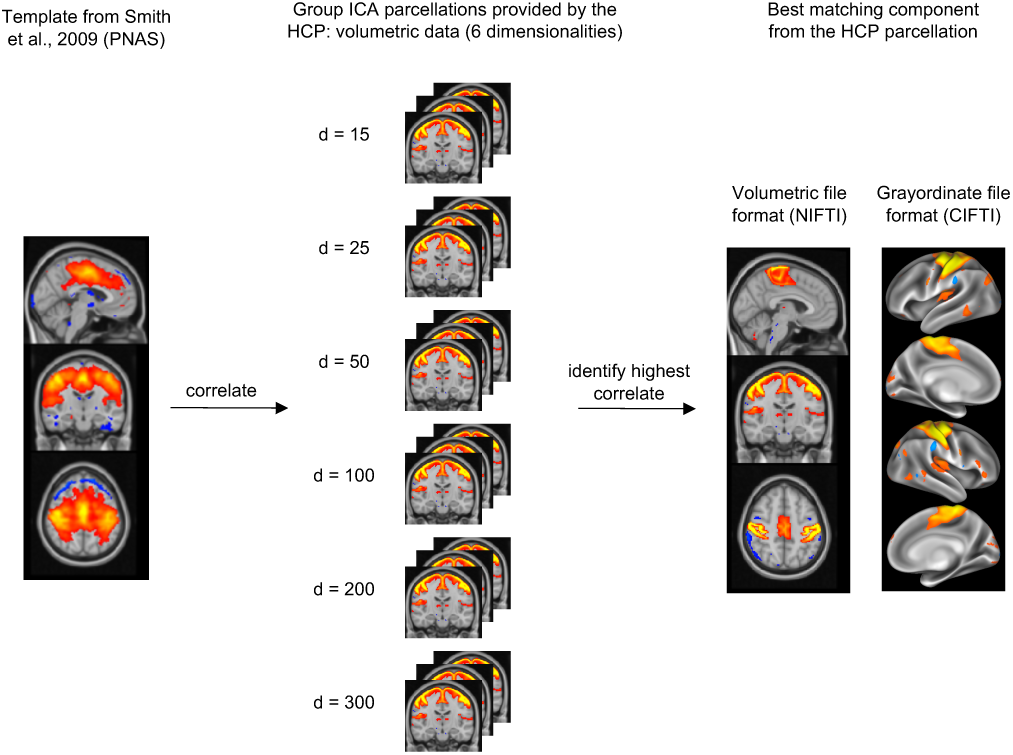
Overview of the selection process. For each of the ten templates proposed by Smith et al. (2009), we ran a correlation analysis with the component maps provided by the Human Connectome Project (HCP) using independent component analysis (ICA, middle panel). We ran correlations with all of the parcellations to identify the best-matching component across all dimensionalities (right panel). We selected that HCP-component with the highest correlation value to the original template as “best match”. We ran this analysis in volumetric space, but also saved a version of each “new” template in the CIFTI file-format (right panel), which is also provided by the HCP.

## Methods

### Subjects and preprocessing

A subset of 1003 subjects from the Human Connectome Project 1200 Subjects Release (www.humanconnectome.org) completed four resting-state fMRI runs (15 minutes each, resulting in 4800 total timepoints per subject at a TR of 0.7 s). The subset consists of 534 healthy females and 469 healthy males, with an age range between 22 and 40 years for both females and males. All study procedures of the HCP protocol were approved by the Institutional Review Board at Washington University in St. Louis. All of the data was fully preprocessed (for the preprocessing procedure, see (Fischl, 2012; Glasser et al., 2013; Jenkinson, Bannister, Brady, Smith, 2002; Jenkinson, Beckmann, Behrens, Woolrich, Smith, 2012; Smith et al., 2013), including artifact removal with ICA-FIX (Griffanti et al., 2014; Salimi-Khorshidi et al., 2014). The preprocessed data was used for a group-ICA parcellation in grayordinate space using FSL’s MELODIC tool at six dimensionalities with 15, 25, 50, 100, 200 and 300 components, respectively (Beckmann Smith, 2004; Hyvärinen, 1999). The resulting component maps of that parcellation procedure are also provided within the HCP database (“HCP1200 Parcellation+Timeseries+Netmats (PTN)”). Next to the grayordinate-based IC maps (i.e. “CIFTI”-files), also voxel-based versions (normalized to volumetric MNI152 2mm space) of these maps were generated and released (i.e. “NIFTI”-files). To note, as in any ICA, these maps are not binary; rather they provide weight values of how strongly each grayordinate/voxel belongs to a component. Thus, every grayordinate/voxel is part of each component, to a given extent, and as a consequence, IC maps can overlap.

### Original approach (Smith et al., 2009)

Since our selection of components from the HCP parcellation was based on the templates provided by (Smith et al., 2009), here we briefly outline how those original templates were generated: Ten template components had been derived from an ICA performed on resting-state datasets from 36 healthy participants. The resulting components were then compared to activation maps from BrainMap, a large database of fMRI and PET brain activation studies (Fox Lancaster, 2002; Laird, Lancaster, Fox, 2005). The BrainMap activation maps were compared to the ICA components from the resting-state data and a set of ten well-matched pairs was identified and published. These templates thus represent a set of major brain networks, each of which putatively corresponds to a different function, and the resulting networks were named accordingly, e.g. “medial visual areas”, “sensorimotor network”, or “auditory network” (see Table 1 for all component names). The reader is referred to the original publication for the full description of the methods.

**Table 1.** Overview of the best-matching ICA map from the HCP parcellations to the original templates (from Smith et al., 2009).

| Component label from the Smith et al. (2009) ICA | Best match of the HCP’s 1200 Subject’s Release ICA | Correlation value (Pearson’s r) |
| --- | --- | --- |
| Medial visual areas | 15-component parcellation: IC 2 | 0.5927 |
| Occipital pole | 15-component parcellation: IC 3 | 0.6982 |
| Lateral visual areas | 25-component parcellation: IC 4 | 0.6262 |
| Default Mode Network | 25-component parcellation: IC 2 | 0.5761 |
| Cerebellum | 25-component parcellation: IC 22 | 0.3417 |
| Sensorimotor Network | 25-component parcellation: IC 13 | 0.3846 |
| Auditory Network | 50-component parcellation: IC 38 | 0.4798 |
| Executive control network | 100-component parcellation: IC 26 | 0.4234 |
| Frontoparietal network, right | 15-component parcellation: IC 7 | 0.6249 |
| Frontoparietal network, left | 15-component parcellation: IC 5 | 0.6192 |

### Statistics

To identify the best-matching IC component from the HCP parcellation for each of the ten Smith-templates, we proceeded as follows: we ran a correlation analysis for each of the ten Smith-templates with all of the HCP’s component maps in volumetric MNI 2mm space (i.e. with 15+25+50+100+200+300=790 components). For each of the ten templates, we chose that HCP component with the highest correlation value (Pearson’s r) to the original template as a new version of that template. In Figure 1, we illustrate the procedure of this selection process. For each network, we identified the best matching HCP-component according to this procedure; additionally, we saved the corresponding grayordinate-based component to generate a version of the templates in CIFTI file-format. Correlation-analyses were performed with MATLAB R2018b (The Mathworks, Natick, MA, USA) and the MATLAB-based toolbox CoSMoMVPA (Oosterhof, Connolly, Haxby, Rosa, 2016). We display the results in the next section using FSLeyes for NIFTI-files (FMRIB Software Library image viewer, https://zenodo.org/record/3403671) and the Connectome Workbench Viewer for CIFTI-files (Marcus et al., 2013, 2011).

## Results

We provide an overview of the best-matching HCP-provided component for each of the ten Smith-templates in Table 1. The correlation values ranged between r = 0.3417 (cerebellar network) and r = 0.6982 (occipital pole network) (mean = 0.5367, SD = 0.1205). The visual comparison between the original Smith-templates and the volumetric version of our identified components show spatial agreement between the two atlases (Fig. 2; left columns show the original templates, right columns our updated versions). This indicates that the general concept of the Smith-templates is maintained within our templates. However, it becomes evident in Figure 2 that spatial alignment with the borders of the MNI152 template is enhanced in the updated version. This is especially clear for the cerebellar network and the sensorimotor network. In the cerebellar network, the original template does not cover the lower half of the cerebellum. Moreover, no distinction between gray and white matter is evident; moreover, there is no distinction between the cerebellum and the brainstem. In contrast, the atlas we provide covers both upper and lower halves of the cerebellum and additionally aligns with the gray-/white-matter boundaries and respects the border to the brainstem. Considering the sensorimotor network, the original atlas does not align with the upper boundaries of the MNI atlas in the pre- and postcentral gyrus, and seems to be located within the white rather than the gray matter. The version we provide, however, wraps smoothly around the gray-/white-matter boundary and aligns with the gyri/sulci of the pre- and postcentral gyri. Similar observations are made for the comparison of the default mode, the auditory, the executive control, as well as the two frontoparietal networks. In all of these networks, the updated templates that we provide align with the gray-matter/white-matter as well as the gray-matter/dura-mater-boundary. In Figure 3, we show the version of our new atlas in CIFTI file-format.

**Fig. 2.**
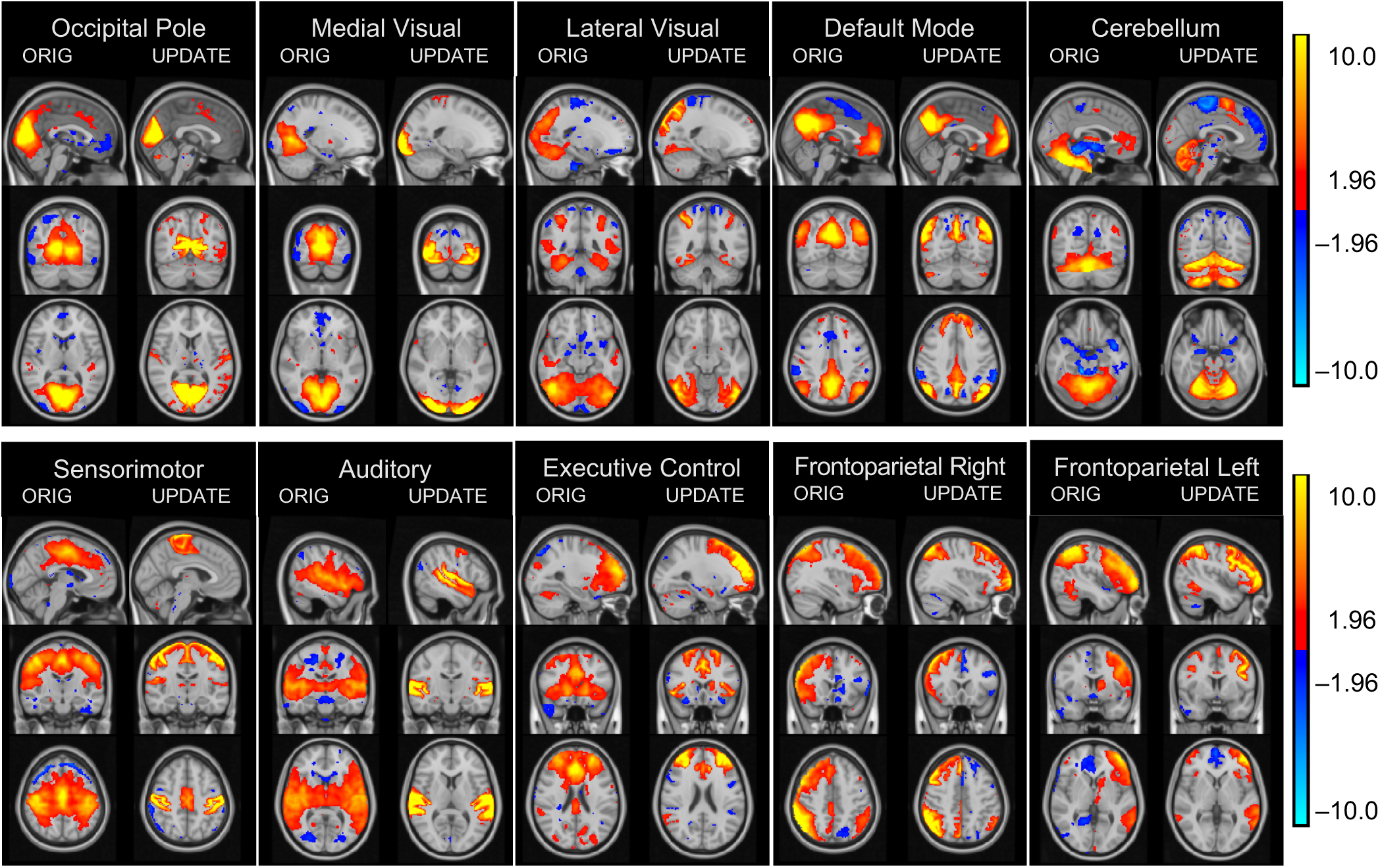
Comparison between the original templates by Smith et al. (2009) (left panels) and the version of those templates generated by the methods we presented (right panels), displayed on the anatomical standard image by the Montreal Neurological Institute (MNI152) in 2mm voxel-resolution. We show z-scored maps of each version to increase comparability. Here it becomes clear that our updated templates are clearly superior to the original ones in terms of alignment with the alignment with the MNI-template as well as the distinction between gray-/white matter boundaries. [Display range: z=1.96–10, negative and positive].

**Fig. 3.**
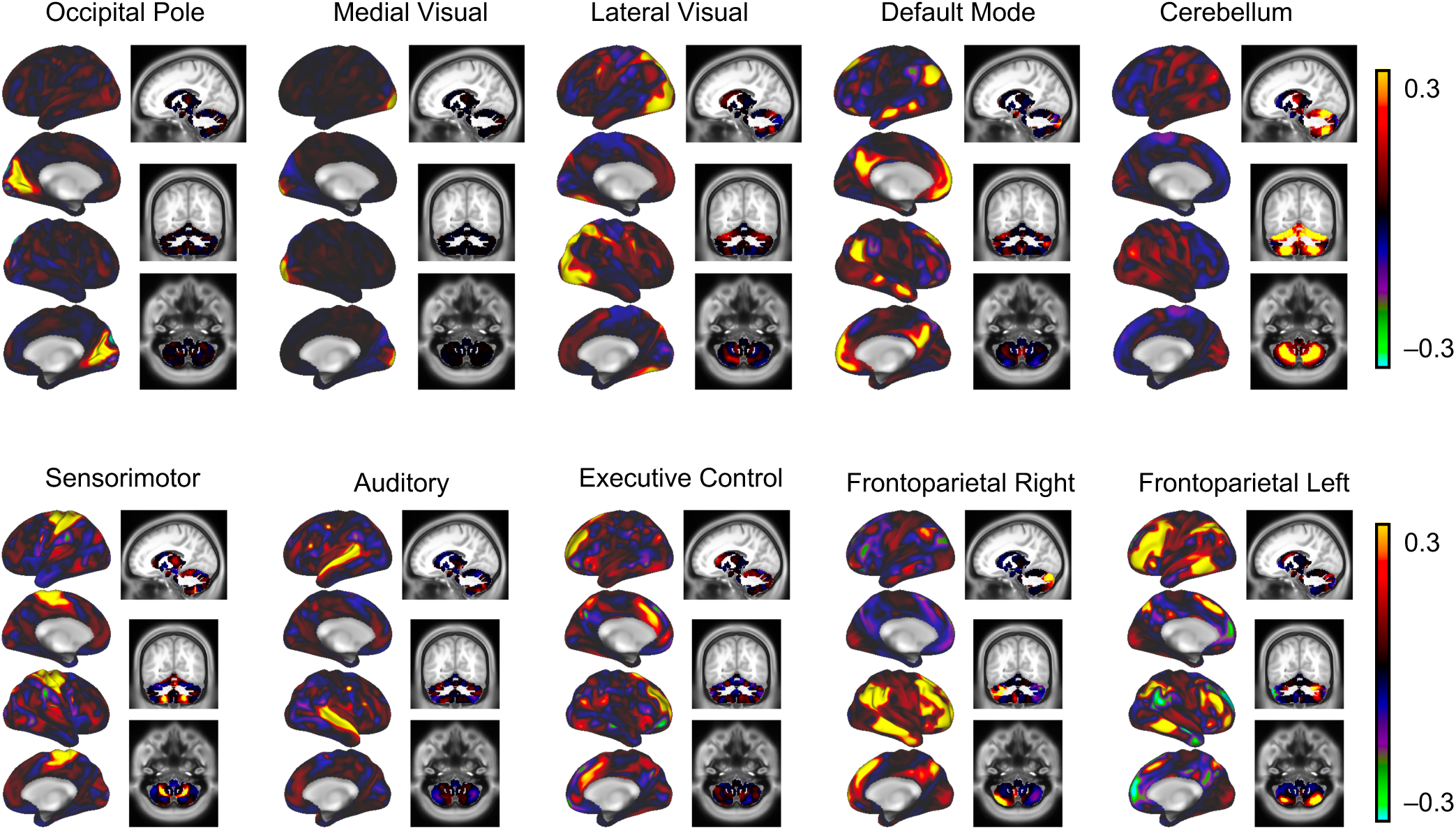
Overview of our ten updated templates in CIFTI file-format, rendered on the surface (left columns) and in volumetric view (right columns, voxel location at –13, –60, –50. [Display range: 0–0.3, negative and positive].

**Fig. 4.**
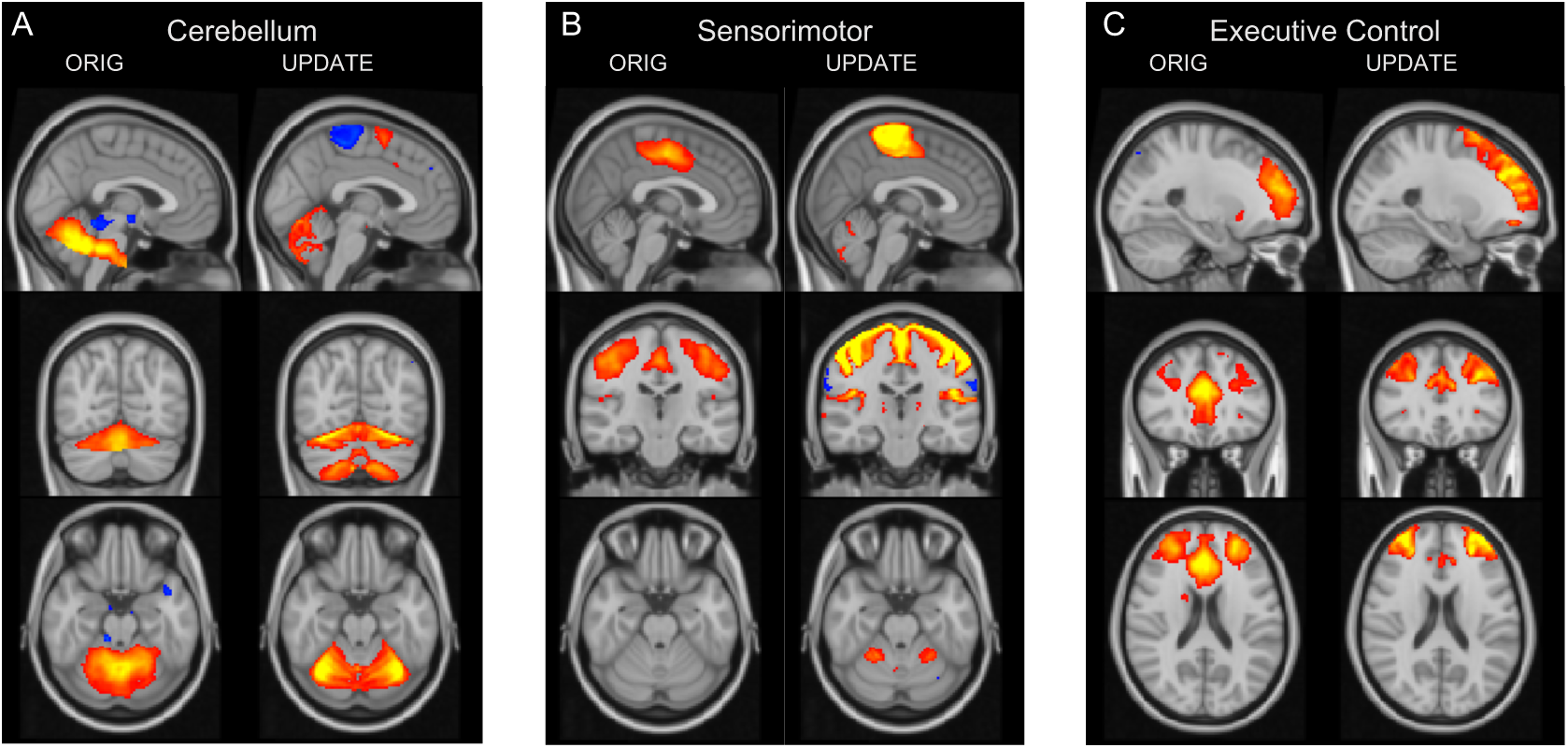
Cerebellar network (A), sensorimotor network (B), and executive control network (C), comparison between the original (left columns) and the updated version (right columns), both displayed on the MNI152 template (2mm). Note the improvements in the updated version in terms of alignment with anatomical boundaries with respect to the gray-/white matter boundaries. Voxel locations: (A) 46, 29, 25; (B) 44, 52, 24; (C) 57, 78, 46. Display ranges: 4-10 for the original templates; 30-100 for the updated versions.

## Discussion

With this work, we have provided a set of ten ICA templates which can be used for group analyses of functional connectivity. This template constitutes a technical improvement of a previously published template (Smith et al., 2009). The original templates offered a consistent set of spatial component maps that additionally can be linked to statistical parameter maps from task-based studies. In our version of the templates, we aimed to improve upon these strengths and therefore do not attempt to take credit for the conceptual aspects of that approach. Instead, we consider our update a major improvement in terms of alignment to anatomical templates (see Fig. 2 for a comparison). The correlation values between the original and our templates (Table 1) support the notion that we succeeded in maintaining the general pattern of the original templates (mean Pearson’s r = 0.5367). However, the maximum correlation between any of the two versions does not exceed 0.7. There are three networks which show especially low correspondence between the two versions, namely the cerebellar (r = 0. 3417), the sensorimotor (r = 0.3846) and the executive control network (r = 0.4234). We argue that the reason for these lower correlations does not indicate conceptual differences between the original templates and our templates but rather a consequence of better spatial alignment of our templates with the anatomical template compared to the original ones. In Figures 4–6, we illustrate the major differences between the templates. Figure 4A shows the comparison between the two cerebellar networks: The original template (Fig. 4A, left column) does not cover the lower half of the cerebellum and does not distinguish between white and gray matter, moreover it spreads into the pons and ventrally to that into cerebrospinal fluid (CSF). The enhanced version (Fig. 4A, right column) covers both halves of the cerebellum and aligns precisely with its lower boundaries, it respects the gray-/white matter boundaries as it wraps around the cerebellum’s foliae, and there are no regions in the pons or outside of the brain covered. Additionally, compared to the original template, the template we provide of the cerebellar network includes also a region of the motor cortex (Fig. 4A, right column, sagittal slice) reflecting the well-known structural and functional connectivity between cerebellum and motor cortex (Daskalakis et al., 2004). The sensorimotor network in the original version (Fig. 4B, left column) is located in the middle of the white matter across the border of the frontal/parietal lobes and is blurred across gray- /white matter and sulci/gyri. In the updated version (Fig. 4B, right column)it aligns well with all these anatomical distinctions and it includes part of the cerebellum (see axial slices of the two versions in Fig. 4B). With respect to the executive control network, Figure 4C illustrates that the updated version (Fig. 4C, left column) better covers the (orbito-) frontal cortex compared to the original version (Fig. 4C, right column). Otherwise, the two versions seem to correspond well in terms of topography. Therefore, sensitivity of FC analyses based on the updated template may profit in the cerebellum, sensorimotor as well as in orbitofrontal cortex, regions that are associated with higher cognitive/motor functions as well as learning. Additionally, we provide a CIFTI-based version of the template, with which we intend to facilitate the inclusion of subcortical structures and the cerebellum in network-based group comparisons on the surface that try to relate to the task-based literature. We believe that one of the applications that will benefit the most from these improvements are dual regression analyses introduced above. To conclude, our update of the ICA-based set of FC templates put forth by Smith et al. (2009) will enhance power, sensitivity, and interpretability of studies interested in testing coherence changes in FC networks.

## ACKNOWLEDGEMENTS

The authors would like to thank Seth M. Levine for his thoughtful comments and efforts toward improving this manuscript.

## Supplementary Material

We provide our template for download in the online supplementary material in both NIFTI and CIFTI file-format at https://github.com/martahedl/Updated-atlas-for-corresponding-brain-activation-during-task-and-rest/tree/master.

## References

Beckmann, C. F., Mackay, C. E., Filippini, N., Smith, S. M. (2009). Group comparison of resting-state FMRI data using multi-subject ICA and dual regression. In OHBM.

Beckmann, C. F., Smith, S. M. (2004). Probabilistic Independent Component Analysis for Functional Magnetic Resonance Imaging. IEEE Transactions on Medical Imaging, 23(2), 137–152. https://doi.org/10.1109/TMI.2003.822821

Biswal, B., Yetkin, F. Z., Haughton, V. M., Hyde, J. S. (1995). Functional connectivity in the motor cortex of resting human brain using echo-planar MRI. Magnetic Resonance in Medicine, 34(4), 537–541. https://doi.org/10.1002/mrm.1910340409

Calhoun, V. D., Liu, J., Adali, T. (2009). A review of group ICA for fMRI data and ICA for joint inference of imaging, genetic, and ERP data. NeuroImage, 45(1Suppl), S163–S172. https://doi.org/10.1016/j.neuroimage.2008.10.057

Castellazzi, G., Debernard, L., Melzer, T. R., Dalrymple-Alford, J. C., D’Angelo, E., Miller, D. H., … Mason, D. F. (2018). Functional Connectivity Alterations Reveal Complex Mechanisms Based on Clinical and Radiological Status in Mild Relapsing Remitting Multiple Sclerosis. Frontiers in Neurology, 9, 690. https://doi.org/10.3389/fneur.2018.00690

Damoiseaux, J. S., Rombouts, S. A. R. B., Barkhof, F., Scheltens, P., Stam, C. J., Smith, S. M., Beckmann, C. F. (2006). Consistent resting-state networks across healthy subjects. Proceedings of the National Academy of Sciences of the United States of America, 103(37), 13848–13853. https://doi.org/10.1073/pnas.0601417103

Daskalakis, Z. J., Paradiso, G. O., Christensen, B. K., Fitzgerald, P. B., Gunraj, C., Chen, R. (2004). Exploring the connectivity between the cerebellum and motor cortex in humans. The Journal of Physiology, 557(Pt 2), 689–700. https://doi.org/10.1113/jphysiol.2003.059808

Fischl, B. (2012). FreeSurfer. NeuroImage, 62(2), 774–781. https://doi.org/10.1016/j.neuroimage.2012.01.021

Fox, M. D., Raichle, M. E. (2007). Spontaneous fluctuations in brain activity observed with functional magnetic resonance imaging. Nature Reviews Neuroscience, 8(9), 700–711. https://doi.org/10.1038/nrn2201

Fox, P. T., Lancaster, J. L. (2002). Opinion: Mapping context and content: the BrainMap model. Nature Reviews. Neuroscience, 3(4), 319–321. https://doi.org/10.1038/nrn789

Glasser, M. F., Sotiropoulos, S. N., Wilson, J. A., Coalson, T. S., Fischl, B., Andersson, J. L., … Hcp, W. (2013). The minimal preprocessing pipelines for the Human Connectome Project. NeuroImage, 80, 105–124. https://doi.org/10.1016/j.neuroimage.2013.04.127

Griffanti, L., Salimi-Khorshidi, G., Beckmann, C. F., Auerbach, E. J., Douaud, G., Sexton, C. E., … Smith, S. M. (2014). ICA-based artefact removal and accelerated fMRI acquisition for improved resting state network imaging. NeuroImage, 95, 232–247. https://doi.org/10.1016/j.neuroimage.2014.03.034

Hyvärinen, A. (1999). Fast and robust fixed-point algorithms for independent component analysis. IEEE Transactions on Neural Networks, 10(3), 626–634. https://doi.org/10.1109/72.761722

Jenkinson, M., Bannister, P., Brady, M., Smith, S. (2002). Improved optimization for the robust and accurate linear registration and motion correction of brain images. NeuroImage, 17(2), 825–841. Retrieved from http://www.ncbi.nlm.nih.gov/pubmed/12377157

Jenkinson, M., Beckmann, C. F., Behrens, T. E. J., Woolrich, M. W., Smith, S. M. (2012). FSL. NeuroImage, 62(2), 782–790. https://doi.org/10.1016/j.neuroimage.2011.09.015

Laird, A. R., Lancaster, J. L., Fox, P. T. (2005). BrainMap: the social evolution of a human brain mapping database. Neuroinformatics, 3(1), 65–78.

Marcus, D. S., Harms, M. P., Snyder, A. Z., Jenkinson, M., Wilson, J. A., Glasser, M. F., … Van Essen, D. C. (2013). Human Connectome Project informatics: Quality control, database services, and data visualization. NeuroImage, 80, 202–219. https://doi.org/10.1016/j.neuroimage.2013.05.077

Marcus, D. S., Harwell, J., Olsen, T., Hodge, M., Glasser, M. F., Prior, F., … Van Essen, D. C. (2011). Informatics and Data Mining Tools and Strategies for the Human Connectome Project. Frontiers in Neuroinformatics, 5(June), 1–12. https://doi.org/10.3389/fninf.2011.00004

Nickerson, L. D., Smith, S. M., Öngür, D., Beckmann, C. F. (2017). Using dual regression to investigate network shape and amplitude in functional connectivity analyses. Frontiers in Neuroscience, 11(MAR), 1–18. https://doi.org/10.3389/fnins.2017.00115

Oosterhof, N. N., Connolly, A. C., Haxby, J. V, Rosa, M. J. (2016). CoSMoMVPA: Multi-Modal Multivariate Pattern Analysis of Neuroimaging Data in Matlab / GNU Octave, 10(July), 1–27. https://doi.org/10.3389/fninf.2016.00027

Pflanz, C. P., Pringle, A., Filippini, N., Warren, M., Gottwald, J., Cowen, P. J., Harmer, C. J. (2015). Effects of seven-day diazepam administration on resting-state functional connectivity in healthy volunteers: A randomized, double-blind study. Psychopharmacology, 232(12), 2139–2147. https://doi.org/10.1007/s00213-014-3844-3

Raichle, M. E., MacLeod, A. M., Snyder, A. Z., Powers, W. J., Gusnard, D. A., Shulman, G. L. (2001). A default mode of brain function. Proceedings of the National Academy of Sciences of the United States of America, 98(2), 676–682. https://doi.org/10.1073/pnas.98.2.676

Rane, S., Mason, E., Hussey, E., Gore, J., Ally, B. A., Donahue, M. J. (2014). The effect of echo time and post-processing procedure on blood oxygenation level-dependent (BOLD) functional connectivity analysis. NeuroImage, 95, 39–47. https://doi.org/10.1016/j.neuroimage.2014.03.055

Rubin, T. N., Koyejo, O., Gorgolewski, K. J., Jones, M. N., Poldrack, R. A., Yarkoni, T. (2017). Decoding brain activity using a large-scale probabilistic functional-anatomical atlas of human cognition. PLoS Computational Biology, 13(10), 1–24. https://doi.org/10.1371/journal.pcbi.1005649

Salimi-Khorshidi, G., Douaud, G., Beckmann, C. F., Glasser, M. F., Griffanti, L., Smith, S. M. (2014). Automatic denoising of functional MRI data: Combining independent component analysis and hierarchical fusion of classifiers. NeuroImage, 90, 449–468. https://doi.org/10.1016/j.neuroimage.2013.11.046

Smith, S. M., Beckmann, C. F., Andersson, J., Auerbach, E. J., Bijsterbosch, J., Douaud, G., … Glasser, M. F. (2013). Resting-state fMRI in the Human Connectome Project. NeuroImage, 80, 144–168. https://doi.org/10.1016/j.neuroimage.2013.05.039

Smith, S. M., Fox, P. T., Miller, K. L., Glahn, D. C., Fox, P. M., Mackay, C. E., … Beckmann, C. F. (2009). Correspondence of the brain’s functional architecture during activation and rest. Proceedings of the National Academy of Sciences of the United States of America, 106(31), 13040–13045. https://doi.org/10.1073/pnas.0905267106

Van Essen, D. C., Smith, S. M., Barch, D. M., Behrens, T. E. J., Yacoub, E., Ugurbil, K., WU-Minn HCP Consortium. (2013). The WU-Minn Human Connectome Project: an overview. NeuroImage, 80(5), 62–79. https://doi.org/10.1016/j.neuroimage.2013.05.041

Yeo, B. T. T., Krienen, F. M., Sepulcre, J., Sabuncu, M. R., others. (2011). The organization of the human cerebral cortex estimated by intrinsic functional connectivity. J Neurophysio, 106, 1125–1165. https://doi.org/jn.00338.2011[pii]10.1152/jn.00338.2011

